# White-matter controllability at birth predicts social engagement and language outcomes in toddlerhood

**DOI:** 10.1101/2025.02.19.638284

**Authors:** Huili Sun, Angelina Vernetti, Marisa Spann, Katarzyna Chawarska, Laura Ment, Dustin Scheinost

## Abstract

Social engagement and language are connected through early development. Alterations in their development can have a prolonged impact on children’s lives. However, the role of white matter at birth in this ongoing connection is less well-known. Here, we investigate how white matter at birth jointly supports social engagement and language outcomes in 642 infants. We use edge-centric network control theory to quantify edge controllability, or the ability of white-matter connections to drive transitions between diverse brain states, at 1 month. Next, we used connectome-based predictive modeling (CPM) to predict the Quantitative Checklist for Autism in Toddlers (Q-CHAT) for social engagement risks and the Bayley Scales of Infant and Toddler Development (BSID-III) for language skills at 18 months from edge controllability. We created the social engagement network (SEN) to predict Q-CHAT scores and the language network (LAN) to predict BSID-III scores. The SEN and LAN were complex, spanning the whole brain. They also significantly overlapped in anatomy and generalized across measures. Controllability in the SEN at 1 month partially mediated associations between Q-CHAT and BSID-III language scores at 18 months. Further, controllability in the SEN significantly differed between term and preterm infants and predicted Q-CHAT scores in an external sample of preterm infants. Together, our results suggest that the intertwined nature of social engagement and language development is rooted in an infant’s white-matter controllability.

**Significance Statement:** During infancy and toddlerhood, social engagement and language emerge together. Delays are often observed in both simultaneously. These interactions persist into later childhood, potentially affecting life quality. We reveal that the interplay between social engagement and language milestones in toddlerhood is rooted in the infant’s structural connectivity, which may assist in early risk identification of developmental delays. Insights into the early brain foundations for emerging social engagement and language skills may open opportunities for individualized interventions to improve developmental outcomes.

## Introduction

From birth to two, infants reach developmental milestones across multiple domains, including sensory-motor^1^, language^2^, attention^3^, and social engagement^4^. Despite occurring in different domains, these milestones emerge close temporally and interact. Further, delays are often observed in multiple domains simultaneously. Particularly robust interactions exist between the social engagement and language domains^5–8^. Social engagement, including gestural communication (e.g., pointing), facilitates word learning^9,10^. Moreover, pointing delays are diagnostic for language deficits^11^, and pointing is a viable target of language intervention^12^.

Similarly, language is central to social cognition and allows for social understanding^13^. Preschool children with language deficits demonstrate bidirectional associations between social and communicative challenges, suggesting that language acquisition fosters adaptive social skills^14^. Further, interactions between social engagement and language persist into later childhood, potentially affecting life quality^15^. For example, both are affected in autism spectrum disorders^16^.

White matter experiences rapid and dynamic changes during infancy, providing the foundation for later milestones^17^. Myelination starts prenatally, enables efficient connectivity between brain regions, and correlates with language and social engagement in toddlerhood^18,19^. For example, the amygdala—a key region for social processing—goes through prominent perinatal myelination, supporting future functional maturation^20^. Similarly, most white-matter tracts are present at birth, forming the structural connectome^3,7^. An infant’s structural connectome is highly organized^21^, correlating with later language and social abilities^22–24^. However, several gaps exist. First, most studies employ correlation or regression rather than predictive models. Predictive models are designed to protect against overfitting, increasing the likelihood of generalization in novel samples. Second, they use standard measures of structural connectivity. Cutting-edge computation models offer information beyond these standard measures. Last, they examine social engagement and language outcomes independently, missing potential interactions between social engagement and language development.

Combining edge-centric network control theory (E-NCT) and connectome-based predictive modeling (CPM) provides a powerful framework to address these gaps. NCT quantifies controllability, or the ability of gray matter regions to drive transitions between diverse brain states^25^. E-NCT extends traditional NCT to an edge-centric view, quantifying the controllability of white matter connections^26^. CPM is a well-established, data-driven method for identifying complex networks subserving multifaceted behaviors (like the interplay between social engagement and language outcomes). Moving from node to edge controllability allows CPM to be applied to controllability data, improving the prediction of phenotypic information over node controllability. While emerging works have started to study infant brain development with NCT^27^ and CPM^22,28^, characterizing edge controllability in infants is novel.

We quantified edge controllability for 642 infants from the developing Human Connectome Project (dHCP)^29^. Using CPM, we defined data-driven networks that predict social engagement and language outcomes at 18 months in term infants. Next, we investigated the interactions between these predictive networks and generalized them to preterm infants. Finally, we performed sensitivity analyses demonstrating that traditional structural connectivity cannot uncover these associations. Our results reveal complex interactions between social engagement and language milestones in the developing brain and may assist with identifying early risk for developmental delays.

## Results

Using E-NCT, we calculated the edge controllability of the structural connectomes for 642 infants (448 term and 192 preterm infants; Table 1) from the dHCP. We examined its associations with long-term language and social engagement outcomes (Fig.1). Diffusion-weighted imaging data was reconstructed using generalized q-sampling imaging. We created structural connectomes for each infant with mean quantitative anisotropy value for each brain region in an infant-specific atlas with 90 nodes. Edge average controllability was then calculated from the structural connectome for each neonate. Developmental outcomes at 18 months corrected age were measured with the Quantitative Checklist for Autism in Toddlers^30^ (Q-CHAT) for social-engagement risks and the Bayley Scales of Infant and Toddler Development^31^ (BSID-III) for language skills. We removed five items from the Q-CHAT related to language to avoid overlap between the developmental outcomes (labeled reduced Q-CHAT). We used CPM^32^ to predict the neurodevelopmental outcome scores for each infant from edge controllability.

**Fig. 1.**
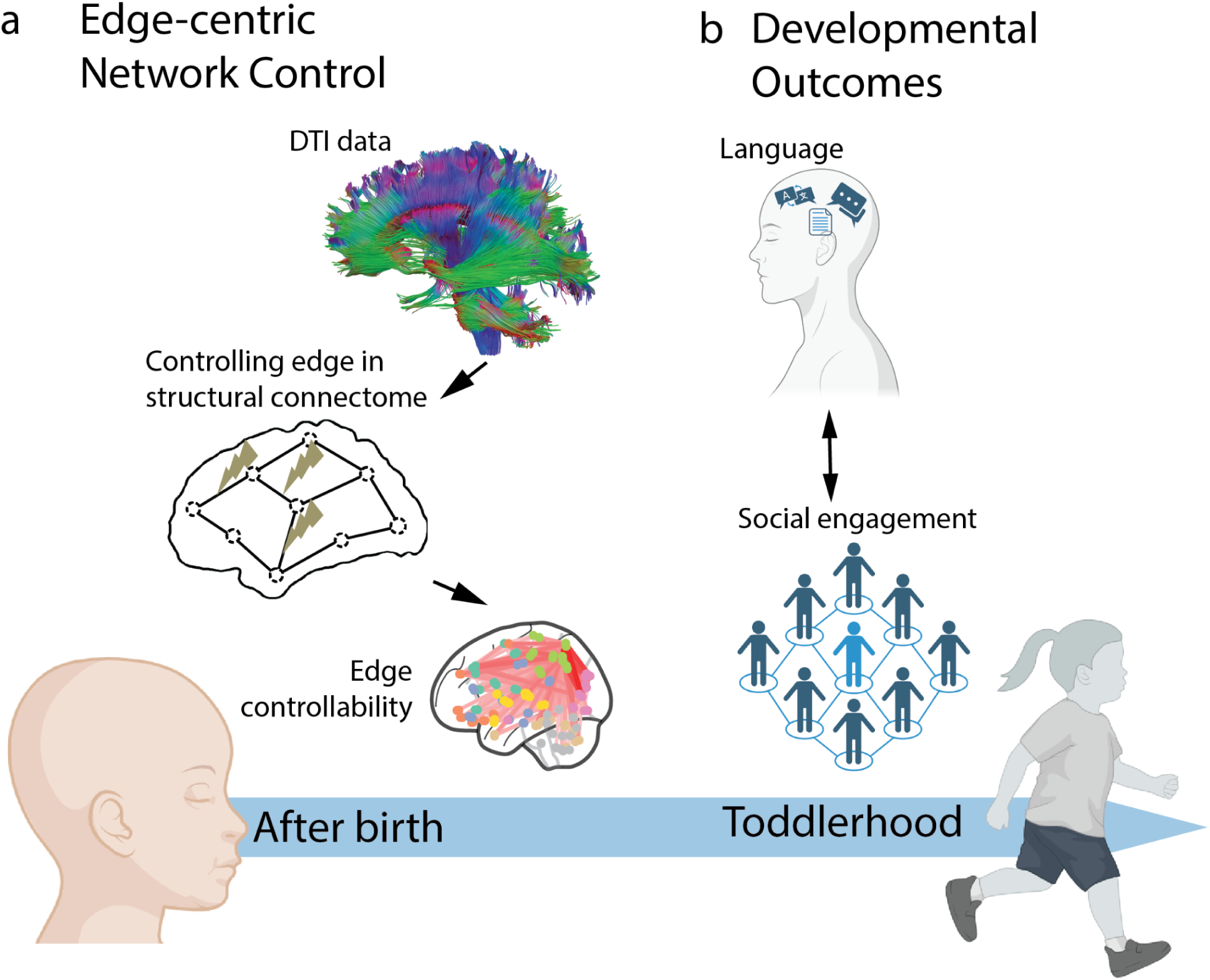
Overview. Structural connectomes were created for 642 infants scanned after birth using diffusion tensor imaging data. **Edge controllability** (a) was calculated from structural connectomes to quantify the capacity of white-matter connections to regulate transitions between diverse brain states. Using connectome-based predictive modeling (CPM), edge controllability during infancy was used to map **brain-behavioral association** (b) with language and social engagement outcomes during toddlerhood.

**Table 1.** Demographics.

| Mean(std) |  | Preterm<br>n=194 | Term n=448 | *t* | *p* |
| --- | --- | --- | --- | --- | --- |
| Gestational Age (GA) at birth/weeks |  | 32.01(3.70) | 39.93(1.26) | -40.27 | 1.29E-177 |
| Postmenstrual Age (PMA) at scan/weeks |  | 35.47(3.60) | 41.19(1.72) | -27.24 | 4.54E-109 |
| BSID-III | Cognition Composite | 98.98(12.56) | 100.52(10.62) | -1.4 | 0.16 |
|  | Language Composite | 96.57(17.29) | 98.07(14.96) | -0.97 | 0.33 |
|  | Motor Composite | 99.54(11.07) | 102.06(9.59) | -2.56 | 0.01 |
| Q-CHAT | Total scores | 30.47(10.38) | 30.04(8.78) | 0.46 | 0.64 |
|  | Reduced scores | 22.88(11.09) | 22.46(9.14) | 0.44 | 0.66 |
|  | > clinical cut-off (%) | 15.98 | 14.06 | 0.63 | 0.53 |

### Edge controllability at birth predicts social engagement in toddlerhood

Using edge controllability and CPM, the reduced Q-CHAT scores were accurately predicted for term infants (n=348; r=0.21, p=1.07e-4; MAE=7.26; Fig. 2B). 170 edges were extracted to form a social engagement network (SEN) to predict reduced Q-CHAT scores for term infants. 116 edges positively correlated with the reduced Q-CHAT scores, connecting mainly the visual and somatomotor, dorsal attention, and subcortical networks, especially among left orbitofrontal inferior, olfactory, inferior occipital lobes, fusiform gyrus, and precuneus. 54 negatively-correlated edges concentrated within subcortical networks and between the subcortical network and somatomotor, ventral attention networks, including the middle cingulate gyrus and calcarine cortex (Fig. 2a).

**Fig. 2.**
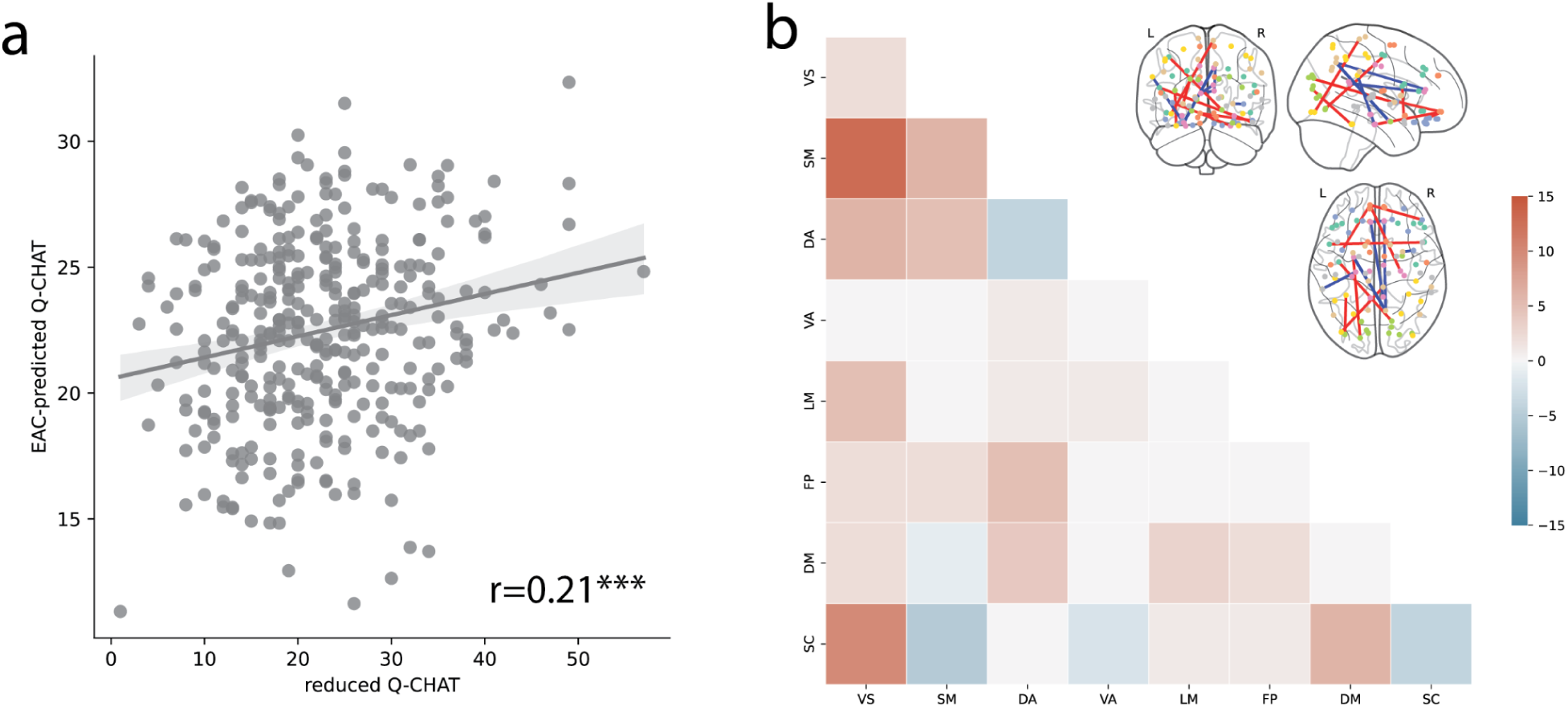
**(A) Predicted reduced Q-CHAT scores** were significantly correlated with the observed reduced Q-CHAT scores (r=0.21, p=1.07e-4)**. (B) Edge controllability** of the social engagement network (SEN) predicts social engagement in term infants. Each element shows the number of edges between each pair of canonical networks in the SEN. (VS visual, SM somatomotor, DA dorsal attention, VA ventral attention, LM limbic, FP frontoparietal, DM default mode, SC subcortical).

### Edge controllability at birth predicts language abilities in toddlerhood

Similar to the prediction of social engagement outcomes, BSID-III language scores were accurately predicted for term infants (n=356; r=0.25, p=2.62e-6; MAE=3.95; Fig. 3a). 344 edges were extracted to form a language abilities network (LAN) for term infants. 128 positively related to the BSID-III language scores, mainly linking the somatomotor, frontoparietal, and default mode networks, especially among precentral, orbitofrontal inferior lobes and rolandic operculum. 216 edges correlated negatively, mainly connecting the visual and subcortical networks between the superior frontal dorsal, inferior frontal opercular, and inferior occipital lobes (Fig. 3b).

**Fig. 3.**
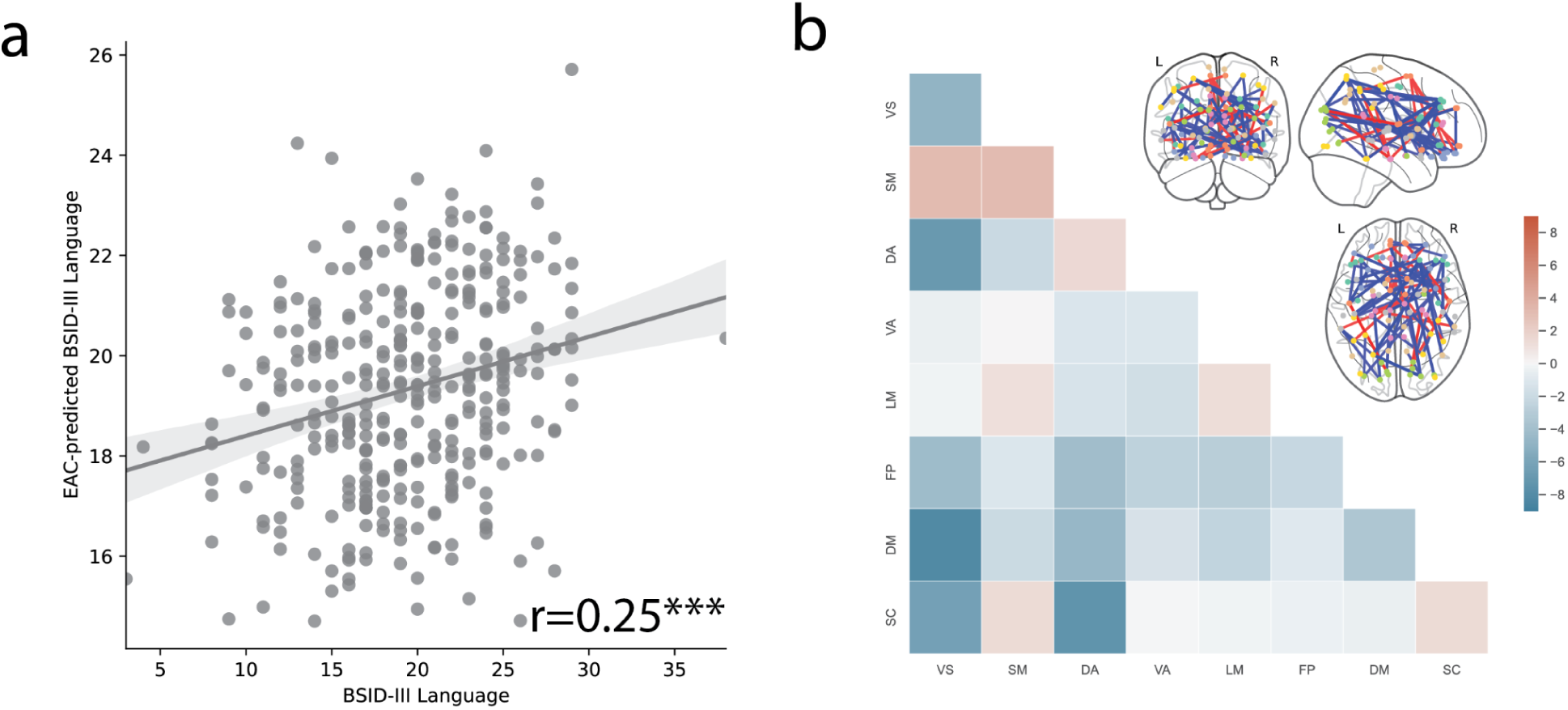
**(A) Predicted BSID-III language scores** were significantly correlated with the observed BSID-III language scaled scores (r=0.25, p=2.62e-6)**. (B) Edge controllability of the language network (LAN)** predicts language outcomes at 18 months in term infants. Each element shows the number of edges between each pair of canonical networks in the LAN. (VS visual, SM somatomotor, DA dorsal attention, VA ventral attention, LM limbic, FP frontoparietal, DM default mode, SC subcortical).

**Fig. 4.**
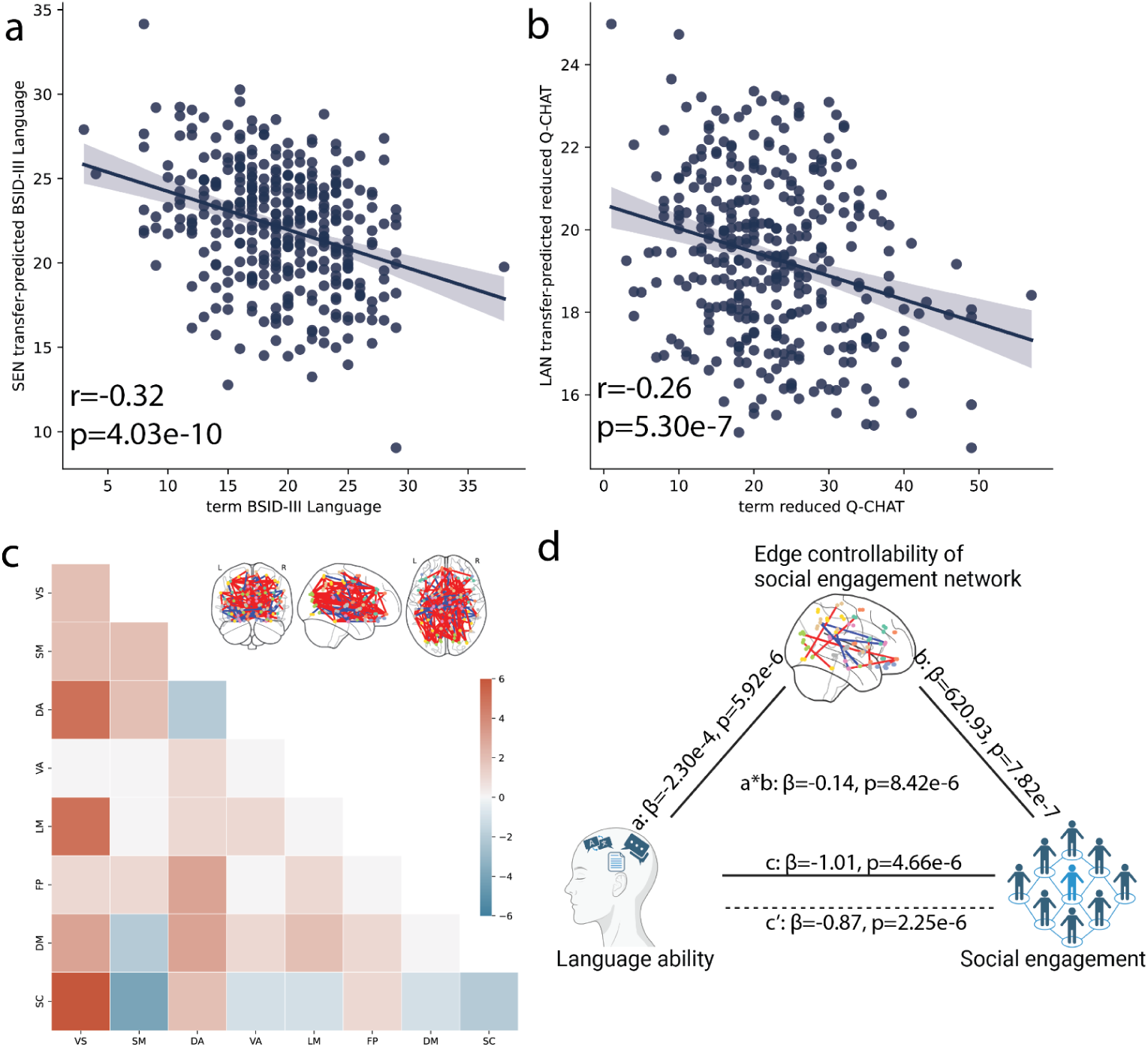
Cross-prediction between LAN and SEN. **(a)** The pre-trained SEN model predicts the language outcomes (r=-0.32, p=4.03e-10). **(b)** The pre-trained LAN model predicts social engagement (r=-0.26, p=5.30e-7). **(c)** Overlap between LAN and SEN **(d)** Edge average controllability in SEN at birth mediated the association between language and social engagement at 18 months. (VS visual, SM somatomotor, DA dorsal attention, VA ventral attention, LM limbic, FP frontoparietal, DM default mode, SC subcortical).

### Interaction between the SEN and LAN

We investigated the interplay between the SEN and LAN. First, we tested the spatial overlap of the SEN and LAN. On the network level, SEN and LAN have nearly 50% overlap in brain connections, involving 58 positive and 25 negative edges (p<0.001). Overlapping positive edges were mainly between the visual network and the dorsal attention, default mode, and subcortical networks. Overlapping negative edges were between the somatomotor and subcortical networks. Second, we tested if the SEN predicted language outcomes and if the LAN predicted social engagement outcomes. Edge average controllability in the SEN predicted the language scores (r=-0.32, p=4.03e-10). The predicted scores are inversely correlated with language, consistent with the negative correlation between the reduced Q-chat and BSID-III language scores (r=-0.63, p=7.93e-54). Similarly, edge average controllability in the LAN predicted the Q-CHAT scores (r=-0.26, p=5.3e-7). Finally, given these associations, we investigate if the SEN or LAN mediated the associations between BSID-III language and the reduced Q-CHAT scores. Using bootstrapped mediation analysis, edge controllability in the SEN partially mediated the association between BSID-III language and the reduced Q-CHAT scores (β=-0.14, p=8.27e-6). Similarly, edge controllability in the SEN partially mediated the association between the reduced Q-CHAT scores and BSID-III language (β=-0.02, p=0.015).

### Network controllability of the SEN and LAN in preterm infants

We investigated the SEN and LAN in preterm infants, a population at risk of social and language deficits. At term equivalent age, preterm infants exhibited significantly lower edge controllability in the SEN but not the LAN (SEN: t=4.54, p=6.96e-6, df=530; LAN: t=1.37, p=0.08, df=530; Fig. 5a-b). In addition to differences in the SEN, preterm infants exhibited lower edge controllability in several canonical network pairs (Fig. S1; Table S1). Next, we generalized the SEN and LAN generated in term infants to the preterm infants. The SEN accurately predicted the reduced Q-CHAT scores of preterm infants (r=0.32, p=0.0097, MAE=9.06; Fig. 5c). However, the LAN did not generalize to the preterm infants (r=0.026, p=0.84, MAE=5.02).

**Fig. 5.**
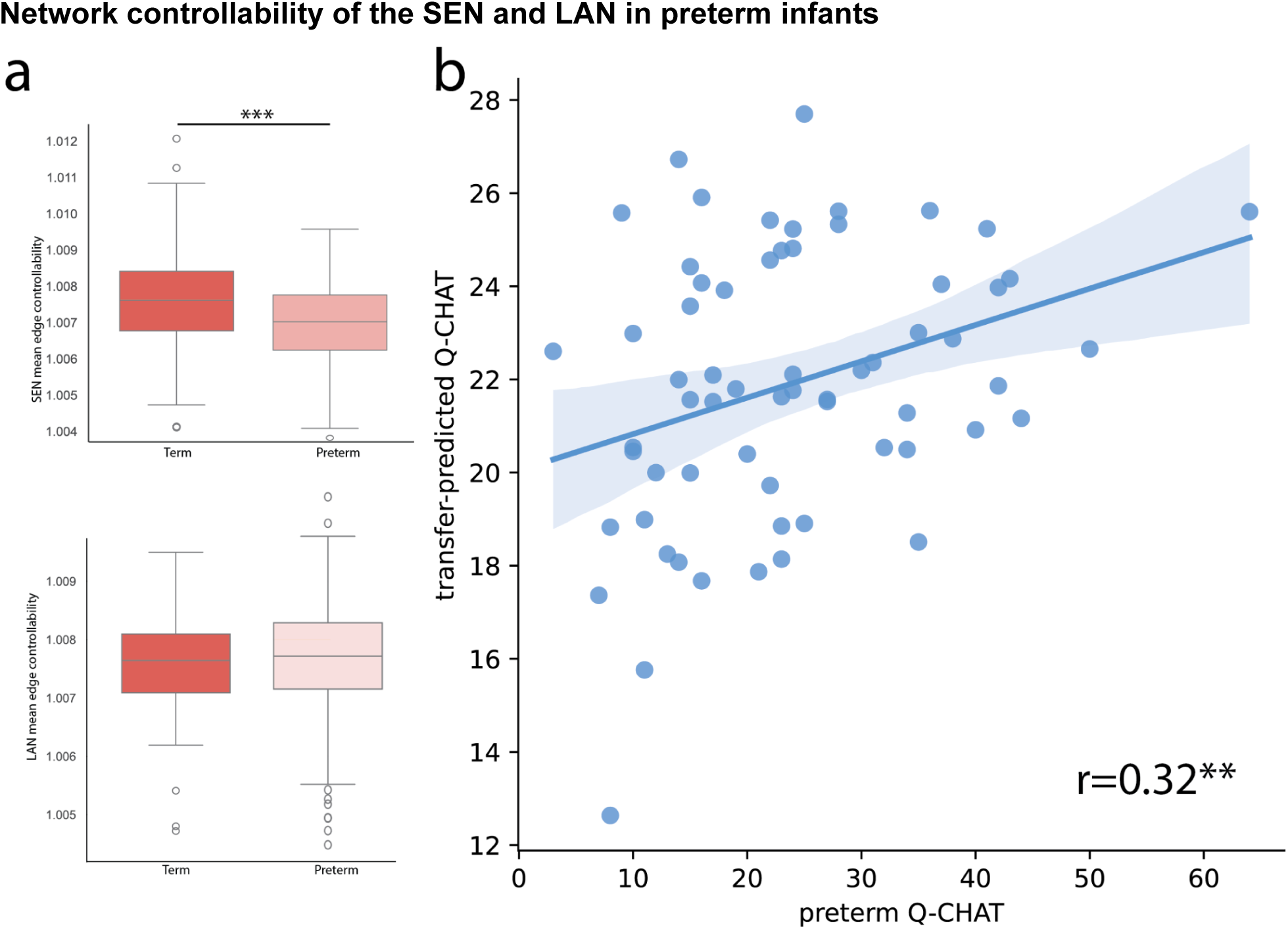
Differences in edge controllability between term and preterm infants. **(a)** Significant group difference between term and preterm infants in edge controllability was observed in SEN but not LAN. **(b)** The SEN, pretrained in term infants, predicted reduced Q-CHAT scores in preterm infants (r=0.32, p=0.0097).

### Sensitivity analysis

We performed a series of sensitivity analyses to ensure the specificity of our results. First, we repeated analyses with the BSID-III cognitive scores given the correlation of the BSID-III language and cognitive scores (r=0.61, p=1.52e-53). BSID-III cognitive scores were accurately predicted for term infants (n=356; r=0.21, p=8.28e-5; MAE=1.64; Fig. S2). Like the LAN, the resulting model also generalized and predicted reduced Q-CHAT scores (Fig. S3). Second, we repeated our analysis using standard structural connectivity rather than edge controllability. CPM with structural connectivity only successfully predicted the reduced Q-CHAT for term infants (r=0.19, p=2.63e-4, MAE=7.40) but not language or cognition scores, aligned with the previous study using the same dataset^28^. This model failed to generalize to the preterm cohort for the reduced Q-CHAT (r=0.24, p=0.054) and to different behavioral domains for term infants (BSID-III language: r=0.0040, p=0.94; BSID-III cognitive scores: r=0.058, p=0.28). These results suggest that edge controllability better captures social engagement and language interactions in the neonatal brain. Finally, we investigate the sensitivity of our results to the items included in our reduced Q-CHAT. First, we repeated analyses using all items in the Q-CHAT. Q-CHAT scores were accurately predicted for term infants (n=348; r=0.22, p=4.32e-5; MAE=6.91). Edge average controllability for that model predicted the language scores (r=-0.30, p=1.37e-8). Second, we removed four other items related to social communication (items 2, 5, 12, and 19). Edge controllability accurately predicted this more constrained Q-CHAT score (r=0.14, p=0.0082), and the model generalized to language outcomes (r=-0.34, p=4.94e-11). These results suggest that social engagement and language interactions are not due to measuring overlapping domains.

## Discussion

We leveraged edge-centric network control theory (E-NCT) and connectome-based predictive modeling (CPM) to predict language and social engagement outcomes in 642 infants. Models significantly predicted reduced Q-CHAT and BSID-III language scores at 18 months using edge controllability at 1 month. The networks that predicted scores were complex, spanning the whole brain. They also significantly overlapped in anatomy and generalized across measures, suggesting that the interaction between social engagement and language development is programmed in the infant’s brain. Controllability in the SEN at 1 month partially mediated associations between reduced Q-CHAT and BSID-III language scores at 18 months. Further, controllability in the SEN significantly differed between term and preterm infants and predicted reduced Q-CHAT scores in an external sample of preterm infants. Finally, sensitivity analyses demonstrate that standard structural connectivity measures could not reveal the same associations, suggesting the value of edge controllability over structural connectivity. In sum, edge controllability at birth shapes toddler behavior and may be a risk marker of future development delays. Moreover, our results reveal a neurobiological pathway for the complex interplay between social engagement and language milestones in early life.

The SEN and LAN overlap with regions and connections of canonical social and language processing circuits in adults. For example, the SEN consists of large areas from the visual and sensorimotor cortices, which play essential roles in social-cognitive tasks^33^. Alterations in these regions are associated with neurodevelopmental disorders, such as autism^34^. The SEN also contains regions related to social skills^35,36^, including the visual, prefrontal, and subcortex.

Similarly, the LAN included Broca’s, Werinicke’s, and other language processing regions^37^. Together, our data-driven models align with previous work in older children and adults, providing initial anatomical validity of our models. Further, they suggest that the foundation of mature adult circuits is present near or even before birth. Notably, the significant overlap in imaging findings mirrors the concordance in the appearance of social behavior and language in the developing brain. Fetuses increase their heart rates to their mother’s voices at 32–34 weeks PMA^38^, exhibit differential responses to their own mother’s voice compared to that of a female stranger reading the same story,^39^ discriminate a change in gender of a speaker reading a sentence^40^, and respond to a prenatal language learning task administered at the beginning of the third trimester^41^. Finally, newborns recognize their mother’s voices and differentiate novel languages on the first postnatal day^42^, providing behavioral data that supports our neuroimaging findings.

However, the SEN and LAN also exhibit striking differences from adult circuits. Language and social processing circuits are highly lateralized in adults^43^, with language processing in the left hemisphere and social processing in the right. Further, homologs of language areas in the right hemisphere process social information. Similarly, lesion studies consistently show altered social processing with right hemisphere lesions, leading to the right-hemispheric dominance hypothesis of social-emotional processing^44^. Broadly, while similar regions process language and social cues in adults (e.g., the inferior frontal gyrus [IFG]), there is hemispheric segregation^43^ (e.g., left IFG for language; right IFG for social skills). Our results suggest widespread, overlapping networks for language and social processing exist during early infancy instead of the hemispheric lateralization observed in adults. Speculatively, this lateralization likely develops later as language and social skills crystallize.

Preterm infants face an increased risk of language and social deficits^45,46^, as well as neurodevelopmental disorders, including ADHD, anxiety, and autism spectrum disorder^47–49^. The preterm group exhibited weaker edge controllability in the SEN. Further, the SEN—trained in term infants—generalized to preterm infants. These results suggest that a similar control network underlies social processing in term and preterm infants. However, preterm infants have lower controllability in these structural connections, speculatively leading to social deficits. In contrast, the LAN was not different between term and preterm infants, possibly secondary to the significant language exposure preterm neonates experience from their first postnatal day and the correlation of this exposure to language outcomes^50,51^. Further, the LAN did not generalize across groups, suggesting a different language network exists in preterms, aligned with extant literature^52,53^. From infancy to adulthood, mounting evidence suggests that those born preterm have an altered language network^54,55^. These alterations have been observed in the fetal period before preterm birth^56^. Overall, E-NCT and the SEN may be markers of future neurobehavioral disorders risk in preterm infants.

Our work has several strengths. First, several recent studies link developmental outcomes and neurodevelopmental risk with infant connectivity data. Multiple brain features correlate with these measures, including small-world topology^57^, morphological^58^, and microstructural features^24^ of structural networks and neural flexibility^59^ and efficiency^60^ of the functional networks. Nevertheless, edge controllability produces more accurate predictive models. E-NCT models brain dynamics on top of the structural connectome, going beyond connectivity and increasing power. Second, toddlers with language delays face higher risks of poor social competence^61^ and behavioral problems^62^. Similarly, toddlers with social deficits may exhibit delayed language skills due to limited social interactions with peers and misinterpreting emotional expression^63^. We traced this connection between language and social processing to right after birth, far earlier than the emergence and observation of behaviors. Finally, we performed several sensitivity analyses and external validations.

Several limitations of our work exist. First, the lack of neuroimaging data between birth and 18 months prevents a complete understanding of brain, language, and social development. Potential sensitive periods might be missed^64,65^. Longitudinal neuroimaging and behavioral data would facilitate creating growth curves—a gold-standard developmental neurobiology approach. Large-scale, longitudinal studies, like the HEALthy Brain and Child Development (HBCD)^66^ Study, may fill this gap. Second, pre- and postnatal environmental factors, such as socioeconomic status and parent-child interactions, affect brain and cognitive development. Similarly, we lack longitudinal, detailed environmental factors in the dHCP cohort. A goal is to identify early risks and provide targeted interventions. In that case, an essential next step is determining which environmental factors mediate models that predict toddler behavior from infant neuroimaging data. Factors associated with better-than-predicted outcomes are strong therapeutic targets. Third, we lacked genetic information. Genetics could assist in determining which and by how much environmental factors can modify predicted outcomes. Fourth, the Q-CHAT was designed to screen for autism and only broadly measures social engagement^28^. Similarly, social engagement is a complex construct with several specific components. Future work is needed to replicate our results with more specific measures of social engagement. Finally, we did not include functional neuroimaging data. While controllability models functional dynamics from structural connectivity, functional data may be more closely associated with later behavior. Task data from awake infant fMRI studies is promising future research for understanding the interplay between social engagement and language development^67,68^.

In conclusion, we leveraged network control theory and machine learning advances to study the interactions between language and social outcomes. Our findings show that controllability after birth can predict both outcomes with overlapping networks. Insights into the early brain foundations for emerging behavior may open opportunities to design individualized interventions for better developmental outcomes.

## Methods

### Participants

Data used here were acquired from the Developing Human Connectome Project (dHCP), a large open science study of infant brain development^69^. Altogether, 642 infants were included in this study, where 448 infants were term infants scanned at 41.19 weeks postmenstrual age (s.d.: 1.72 weeks), 194 were preterm scanned at 35.47 weeks postmenstrual age (s.d.: 3.59 weeks). The study was approved by the National Research Ethics Service West London committee, and written consent was obtained from participating families before imaging. In a follow-up visit at 18 months corrected age, subjects were tested for the developmental outcomes: 420 infants 412 infants (356 terms, 64 preterms) were evaluated with the Bayley Scales of Infant and Toddler Development (BSID-III), and (349 terms, 63 preterms) infants were screened with Quantitative Checklist for Autism in Toddlers (Q-CHAT) to assess the risk in social abilities. We removed items 1, 4, 8, 17, and 18 from the Q-CHAT related to language to avoid overlap between the developmental outcomes (labeled reduced Q-CHAT; Table S2). More demographic information can be found in Table 1.

### MRI Preprocessing

All imaging scans were conducted at the Evelina Newborn Imaging Centre, St Thomas’ Hospital, London, UK, using a Philips Achieva 3 T scanner (Philips Medical Systems, Best, The Netherlands) equipped with a dHCP-customized neonatal imaging system, including a 32-channel receive neonatal head coil (Rapid Biomedical GmbH, Rimpar, DE)^70^. Infants were scanned during unsedated sleep following feeding and immobilization in a vacuum-evacuated bag. They were subjected to hearing protection and physiological monitoring (pulse oximetry, body temperature, and electrocardiography).

T2-weighted images were acquired through a Turbo spin echo sequence with specific parameters (TR=12s, TE=156ms, SENSE factor 2.11 [axial] and 2.54 [sagittal]) with overlapping slices (resolution = 0.8×0.8×1.6 mm^3^) and subsequently motion-corrected and super-resolved to a resolution of 0.8×0.8×0.8 mm^371^. Diffusion-weighted imaging (DWI) was performed in 300 directions (TR=3.8s, TE=90ms, SENSE factor 1.2, multiband factor 4, and resolution 1.5×1.5×3mm^3^ with 1.5 mm slice overlap) with b-values of 400s/mm2, 1000s/mm^2^ and 2600 s/mm^2^ spherically distributed in 64, 88 and 128 directions respectively using interleaved phase encoding^72^.

DWI data underwent reconstruction and correction processes, including denoising, motion correction, eddy current correction, Gibbs ringing correction, and susceptibility artifact correction using the diffusion SHARD pipeline^73^. In-scanner head motion was assessed, and scans with low neighboring DWI correction values were excluded^74^. The accuracy of b-table orientation was verified, and the diffusion data were reconstructed using generalized q-sampling imaging (GQI)^75^. Tensor metrics were calculated using the Extreme Science and Engineering Discovery Environment^76^. Whole-brain fiber tracking was conducted with DSI-studio http://dsi-studio.labsolver.org/), employing quantitative anisotropy (QA) as the termination threshold. QA values were computed in each voxel in their native space for every subject. The tracking parameters were set as the angular cutoff of 60 degrees, step size of 1.0mm, minimum length of 30 mm, and maximum length of 300 mm. The whole-brain fiber tracking process was performed with the FACT algorithm until 1,000,000 streamlines were reconstructed for each individual. The structural connectome for each individual was constructed using neonatal AAL-aligned brain parcellation with 90 nodes^77^. T2-weighted images in native DWI space were used to provide information on region segmentation during the construction of connectomes. The structural connectome for each individual was then constructed with a connectivity threshold of 0.001. Pairwise connectivity strength was calculated as the average QA value of each fiber connecting the two end regions, which results in a 90×90 structural connectome for each participant.

### Edge-centric Network Control Theory (E-NCT)

The E-NCT framework takes two steps^26^. First, to convert from the node-centric view to edge-centric, the adjacency matrix *A*_*N*×*N*_ is converted into edge-centric adjacency matrix 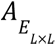 via the incidence matrix *C*^78^. The incidence matrix *C* can be obtained by *CC^T^* = *A* + *D*, where *D* is the node degree matrix with its diagonal elements as 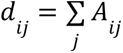. After constructing the incidence matrix, the edge-centric adjacency matrix can be obtained by *A_*E*_* = *C*^*T*^*C* − *W*, where *W* is the diagonal matrix with each element as the weight of each edge. Next, the network control theory^25^ can be applied to the edge-centric adjacent matrix *A_*E*_* to measure the average controllability for edges in the brain network. Average controllability represents an edge’s ability to steer the brain to easily reachable states and is defined as *Trace*(**W**_*L*_), where 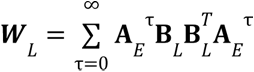.

### Connectome-based Predictive Modeling

We employed connectome-based predictive modeling (CPM) to predict the cognitive developmental outcomes (BSID-III) and social engagement risks (reduced Q-CHAT) for term infants at 18 months with individual edge controllability. The evaluation utilized a 10-fold cross-validation approach, incorporating a feature selection threshold set at p=0.01^32^. All subjects were randomly divided into nine subgroups used for training and the one left for testing. During model training, the related brain features (i.e., edge average controllability) were selected with Pearson’s correlation (p-value < 0.01) within the nine training groups. Next, the selected features for each individual were summarized into a single number. Linear regression was then used to model this summary score and the behavioral scores in the training group. Finally, this model was applied to the testing group. The whole procedure was repeated iteratively, with each subgroup being the testing group once, generating a predicted result for each individual in the dataset. We predicted each behavioral score (i.e., BSID-III, Q-CHAT) independently.

### Network Identification

To identify the social engagement network (SEN) and language network (LAN), CPM for each behavioral score was repeated 100 times. Edges appearing in over 90% of the iterations formed the corresponding network.

## Acknowledgments and funding sources

This study was supported by NIMH (1R01MH137609-01 and 5R01MH126133-02 to D.S.) Data were provided by the developing Human Connectome Project, KCL-Imperial-Oxford Consortium funded by the European Research Council under the European Union Seventh Framework Programme (FP/2007-2013) / ERC Grant Agreement no. [319456]. We are grateful to the families who generously supported this trial.

## Data availability

Raw Data from the Developing Human Connectome Project is publicly available at http://www.developingconnectome.org/data-release/third-data-release and can be downloaded upon request from NDA. Source data are provided in this paper.

## Code availability

Preprocessing code can be found at https://brain.labsolver.org/hcp_d2.html for brain structural connectome. The edge-centric network control theory code is publicly available at https://github.com/huiliii/edge_control.

## Notes

### Competing Interest Statement

The authors have declared no competing interest.

## Reference

1. Arichi, T. et al. Development of BOLD signal hemodynamic responses in the human brain. NeuroImage 63, 663–673 (2012).

2. Dehaene-Lambertz, G., Dehaene, S. & Hertz-Pannier, L. Functional Neuroimaging of Speech Perception in Infants. Science 298, 2013–2015 (2002).

3. Grossmann, T. & Johnson, M. H. Selective prefrontal cortex responses to joint attention in early infancy. Biology Letters 6, 540–543 (2010).

4. Swain, J. E., Lorberbaum, J. P., Kose, S. & Strathearn, L. Brain basis of early parent–infant interactions: psychology, physiology, and in vivo functional neuroimaging studies. Journal of Child Psychology and Psychiatry 48, 262–287 (2007).

5. Beitchman, J. H. et al. Long-Term Consistency in Speech/Language Profiles: II. Behavioral, Emotional, and Social Outcomes. Journal of the American Academy of Child & Adolescent Psychiatry 35, 815–825 (1996).

6. Sundheim, S. T. P. V. & Voeller, K. K. S. Psychiatric Implications of Language Disorders and Learning Disabilities: Risks and Management. J Child Neurol 19, 814–826 (2004).

7. Sahin, N. T., Pinker, S., Cash, S. S., Schomer, D. & Halgren, E. Sequential Processing of Lexical, Grammatical, and Phonological Information Within Broca’s Area. Science 326, 445–449 (2009).

8. Shevell, M. I., Majnemer, A., Webster, R. I., Platt, R. W. & Birnbaum, R. Outcomes at school age of preschool children with developmental language impairment. Pediatric Neurology 32, 264–269 (2005).

9. Booth, A. E., McGregor, K. K. & Rohlfing, K. J. Socio-Pragmatics and Attention: Contributions to Gesturally Guided Word Learning in Toddlers. Language Learning and Development 4, 179–202 (2008).

10. Colonnesi, C., Stams, G. J. J. M., Koster, I. & Noom, M. J. The relation between pointing and language development: A meta-analysis. Developmental Review 30, 352–366 (2010).

11. Lüke, C., Ritterfeld, U., Grimminger, A., Liszkowski, U. & Rohlfing, K. J. Development of Pointing Gestures in Children With Typical and Delayed Language Acquisition. J Speech Lang Hear Res 60, 3185–3197 (2017).

12. LeBarton, E. S., Goldin-Meadow, S. & Raudenbush, S. Experimentally Induced Increases in Early Gesture Lead to Increases in Spoken Vocabulary. Journal of Cognition and Development 16, 199–220 (2015).

13. Kinzler, K. D., Dupoux, E. & Spelke, E. S. The native language of social cognition. Proceedings of the National Academy of Sciences 104, 12577–12580 (2007).

14. Bretherton, L. et al. Developing relationships between language and behaviour in preschool children from the Early Language in Victoria Study: implications for intervention. Emotional and Behavioural Difficulties 19, 7–27 (2014).

15. Van Agt, H., Verhoeven, L., Van Den Brink, G. & De Koning, H. The impact on socio-emotional development and quality of life of language impairment in 8-year-old children. Developmental Medicine & Child Neurology 53, 81–88 (2011).

16. Ingersoll, B., Meyer, K., Bonter, N. & Jelinek, S. A Comparison of Developmental Social–Pragmatic and Naturalistic Behavioral Interventions on Language Use and Social Engagement in Children With Autism. Journal of Speech, Language, and Hearing Research 55, 1301–1313 (2012).

17. Paterson, S. J., Heim, S., Thomas Friedman, J., Choudhury, N. & Benasich, A. A. Development of structure and function in the infant brain: Implications for cognition, language and social behaviour. Neuroscience & Biobehavioral Reviews 30, 1087–1105 (2006).

18. Schneider, N., Greenstreet, E. & Deoni, S. C. L. Connecting inside out: Development of the social brain in infants and toddlers with a focus on myelination as a marker of brain maturation. Child Development 93, 359–371 (2022).

19. O’Muircheartaigh, J. et al. Interactions between White Matter Asymmetry and Language during Neurodevelopment. J. Neurosci. 33, 16170–16177 (2013).

20. Mulc, D. et al. Fetal development of the human amygdala. Journal of Comparative Neurology 532, e25580 (2024).

21. Ball, G. et al. Rich-club organization of the newborn human brain. Proc. Natl. Acad. Sci. U.S.A. 111, 7456–7461 (2014).

22. Sun, H. et al. Brain age prediction and deviations from normative trajectories in the neonatal connectome. Nat Commun 15, 10251 (2024).

23. Paul, L. K., Turner, J., Sung, S. & Elison, J. T. Social and Communication Development in Infants with Isolated Agenesis of the Corpus Callosum. The Journal of Pediatrics: Clinical Practice 14, 200118 (2024).

24. Vaher, K. et al. Neonatal amygdala microstructure and structural connectivity are associated with autistic traits at 2 years of age. 2024.11.29.24318196 Preprint at 10.1101/2024.11.29.24318196 (2024).

25. Gu, S. et al. Controllability of structural brain networks. Nat Commun 6, 8414 (2015).

26. Sun, H. et al. Edge-centric network control on the human brain structural network. Imaging Neuroscience 2, 1–15 (2024).

27. Sun, H. et al. Network controllability of structural connectomes in the neonatal brain. Nat Commun 14, 5820 (2023).

28. Fenchel, D. et al. Neonatal multi-modal cortical profiles predict 18-month developmental outcomes. Developmental Cognitive Neuroscience 54, 101103 (2022).

29. Fitzgibbon, S. P. et al. The developing Human Connectome Project (dHCP) automated resting-state functional processing framework for newborn infants. NeuroImage 223, 117303 (2020).

30. Allison, C. et al. The Q-CHAT (Quantitative CHecklist for Autism in Toddlers): A Normally Distributed Quantitative Measure of Autistic Traits at 18–24 Months of Age: Preliminary Report. J Autism Dev Disord 38, 1414–1425 (2008).

31. Bayley, N. Bayley Scales of Infant and Toddler Development, Third Edition. (2005) doi:10.1037/t14978-000.

32. Shen, X. et al. Using connectome-based predictive modeling to predict individual behavior from brain connectivity. Nat Protoc 12, 506–518 (2017).

33. Grossmann, T. & Johnson, M. H. The development of the social brain in human infancy. European Journal of Neuroscience 25, 909–919 (2007).

34. Ilyka, D., Johnson, M. H. & Lloyd-Fox, S. Infant social interactions and brain development: A systematic review. Neuroscience & Biobehavioral Reviews 130, 448–469 (2021).

35. Nebel, M. B. et al. Intrinsic Visual-Motor Synchrony Correlates With Social Deficits in Autism. Biological Psychiatry 79, 633–641 (2016).

36. Grossmann, T. The role of medial prefrontal cortex in early social cognition. Front. Hum. Neurosci. 7, (2013).

37. Skeide, M. A. & Friederici, A. D. The ontogeny of the cortical language network. Nat Rev Neurosci 17, 323–332 (2016).

38. Kisilevsky, B. S. & Hains, S. M. J. Onset and maturation of fetal heart rate response to the mother’s voice over late gestation. Developmental Science 14, 214–223 (2011).

39. Kisilevsky, B. S. Fetal Auditory Processing: Implications for Language Development? in Fetal Development: Research on Brain and Behavior, Environmental Influences, and Emerging Technologies (eds. Reissland, N. & Kisilevsky, B. S.) 133–152 (Springer International Publishing, Cham, 2016). doi:10.1007/978-3-319-22023-9_8.

40. Danilov, I. V. Ontogenesis of Social Interaction: Review of Studies Relevant to the Fetal Social Behavior. J Med - Clin Res & Rev 4, (2020).

41. Partanen, E. et al. Learning-induced neural plasticity of speech processing before birth. Proc. Natl. Acad. Sci. U.S.A. 110, 15145–15150 (2013).

42. Moon, C., Lagercrantz, H. & Kuhl, P. K. Language experienced in utero affects vowel perception after birth: a two-country study. Acta Paediatrica 102, 156–160 (2013).

43. Rajimehr, R., Firoozi, A., Rafipoor, H., Abbasi, N. & Duncan, J. Complementary hemispheric lateralization of language and social processing in the human brain. Cell Reports 41, 111617 (2022).

44. Palomero-Gallagher, N. & Amunts, K. A short review on emotion processing: a lateralized network of neuronal networks. Brain Struct Funct 227, 673–684 (2022).

45. Allievi, A. G. et al. Maturation of Sensori-Motor Functional Responses in the Preterm Brain. Cerebral Cortex 26, 402–413 (2016).

46. Miller, J. V., Chau, V., Synnes, A., Miller, S. P. & Grunau, R. E. Brain Development and Maternal Behavior in Relation to Cognitive and Language Outcomes in Preterm-Born Children. Biological Psychiatry 92, 663–673 (2022).

47. Bishop, C. L., Lean, R. E., Smyser, T. A., Smyser, C. D. & Rogers, C. E. Adverse Childhood Experiences and Socioemotional Outcomes of Children Born Very Preterm. The Journal of Pediatrics 276, 114377 (2025).

48. Shuster, C. L. et al. Two-Year Autism Risk Screening and 3-Year Developmental Outcomes in Very Preterm Infants. JAMA Pediatr (2023) doi:10.1001/jamapediatrics.2023.5727.

49. Fitzallen, G. C. et al. Risk profiles of the preterm behavioral phenotype in children aged 3 to 18 years. Front. Pediatr. 11, (2023).

50. Caskey, M., Stephens, B., Tucker, R. & Vohr, B. Adult Talk in the NICU With Preterm Infants and Developmental Outcomes. Pediatrics 133, e578–e584 (2014).

51. McGowan, E. C., Caskey, M., Tucker, R. & Vohr, B. R. A Randomized Controlled Trial of a Neonatal Intensive Care Unit Language Intervention for Parents of Preterm Infants and 2-Year Language Outcomes. The Journal of Pediatrics 264, 113740 (2024).

52. Scheinost, D. et al. Preterm birth alters neonatal, functional rich club organization. Brain Struct Funct 221, 3211–3222 (2016).

53. Scheinost, D. et al. Alterations in Anatomical Covariance in the Prematurely Born. Cerebral Cortex 27, 534–543 (2017).

54. Constable, R. T. et al. A left cerebellar pathway mediates language in prematurely-born young adults. NeuroImage 64, 371–378 (2013).

55. Kvanta, H. et al. Language performance and brain volumes, asymmetry, and cortical thickness in children born extremely preterm. Pediatr Res 95, 1070–1079 (2024).

56. Thomason, M. E. et al. Weak functional connectivity in the human fetal brain prior to preterm birth. Sci Rep 7, 39286 (2017).

57. Batalle, D. et al. Altered small-world topology of structural brain networks in infants with intrauterine growth restriction and its association with later neurodevelopmental outcome. NeuroImage 60, 1352–1366 (2012).

58. Fenchel, D. et al. Development of Microstructural and Morphological Cortical Profiles in the Neonatal Brain. Cerebral Cortex 30, 5767–5779 (2020).

59. Yin, W. et al. The emergence of a functionally flexible brain during early infancy. Proceedings of the National Academy of Sciences 117, 23904–23913 (2020).

60. Jang, Y. H. et al. Altered development of structural MRI connectome hubs at near-term age in very and moderately preterm infants. Cerebral Cortex bhac438 (2022) doi:10.1093/cercor/bhac438.

61. Horwitz, S. M. et al. Language Delay in a Community Cohort of Young Children. Journal of the American Academy of Child & Adolescent Psychiatry 42, 932–940 (2003).

62. Rantalainen, K. et al. Early vocabulary development: Relationships with prelinguistic skills and early social-emotional/behavioral problems and competencies. Infant Behavior and Development 62, 101525 (2021).

63. Rautakoski, P. et al. Communication skills predict social-emotional competencies. Journal of Communication Disorders 93, 106138 (2021).

64. Nelson, C. A. & Gabard-Durnam, L. J. Early Adversity and Critical Periods: Neurodevelopmental Consequences of Violating the Expectable Environment. Trends in Neurosciences 43, 133–143 (2020).

65. Margolis, E. T. & Gabard-Durnam, L. J. Prenatal influences on postnatal neuroplasticity: Integrating DOHaD and sensitive/critical period frameworks to understand biological embedding in early development. Infancy **n/a**,.

66. Volkow, N. D. et al. The HEALthy Brain and Child Development Study (HBCD): NIH collaboration to understand the impacts of prenatal and early life experiences on brain development. Developmental Cognitive Neuroscience 69, 101423 (2024).

67. Nielsen, A. N., Graham, A. M. & Sylvester, C. M. Baby Brains at Work: How Task-Based Functional Magnetic Resonance Imaging Can Illuminate the Early Emergence of Psychiatric Risk. Biological Psychiatry 93, 880–892 (2023).

68. Ellis, C. T. & Turk-Browne, N. B. Infant fMRI: A Model System for Cognitive Neuroscience. Trends in Cognitive Sciences 22, 375–387 (2018).

69. Edwards, A. D. et al. The Developing Human Connectome Project Neonatal Data Release. Front. Neurosci. 16, (2022).

70. Hughes, E. J. et al. A dedicated neonatal brain imaging system. Magnetic Resonance in Medicine 78, 794–804 (2017).

71. Cordero-Grande, L., Hughes, E. J., Hutter, J., Price, A. N. & Hajnal, J. V. Three-dimensional motion corrected sensitivity encoding reconstruction for multi-shot multi-slice MRI: Application to neonatal brain imaging. Magnetic Resonance in Medicine 79, 1365–1376 (2018).

72. Scheid, B. H. et al. Time-evolving controllability of effective connectivity networks during seizure progression. Proceedings of the National Academy of Sciences 118, e2006436118 (2021).

73. Christiaens, D. et al. Scattered slice SHARD reconstruction for motion correction in multi-shell diffusion MRI. NeuroImage 225, 117437 (2021).

74. Yeh, F.-C. et al. Differential tractography as a track-based biomarker for neuronal injury. NeuroImage 202, 116131 (2019).

75. Yeh, F.-C., Wedeen, V. J. & Tseng, W.-Y. I. Generalized $ q$-Sampling Imaging. IEEE Transactions on Medical Imaging 29, 1626–1635 (2010).

76. Towns, J. et al. XSEDE: Accelerating Scientific Discovery. Comput. Sci. Eng. 16, 62–74 (2014).

77. Shi, F. et al. Infant Brain Atlases from Neonates to 1- and 2-Year-Olds. PLOS ONE 6, e18746 (2011).

78. Evans, T. S. & Lambiotte, R. Line graphs of weighted networks for overlapping communities. Eur. Phys. J. B 77, 265–272 (2010).

