## Supplementary Information for "White-matter controllability at birth predicts social engagement and language outcomes in toddlerhood"

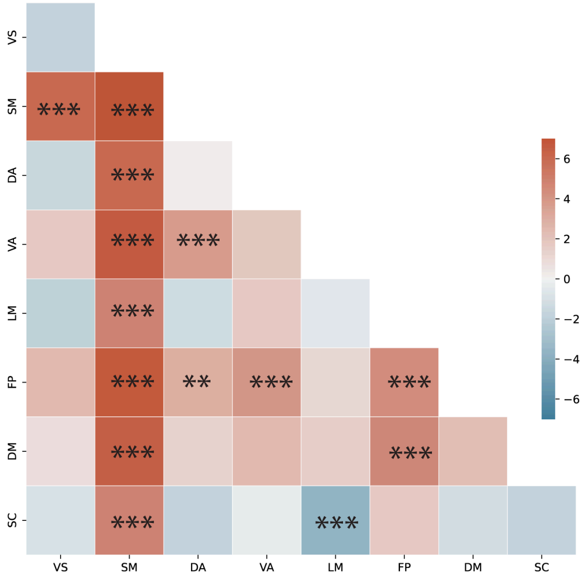


**Fig. S1. Comparisons ‘of edge controllability in canonical network pairs between term and preterm infants.** Over the canonical networks, term infants also showed higher edge average controllability in 13 networks among 36 of them, especially in the visual, somatomotor, and subcortical networks.

Table S1. T-tests of network controllability between preterm and term group (FDR-corrected)

| Network | *t*-value | FDR-*p* value | Network | *t*-value | FDR-*p* value |
|---|---|---|---|---|---|
| VS-VS | 4.73 | 1.12e-5 | FP-DA | 5.63 | 3.44e-7 |
| SM-VS | 4.63 | 1.63e-5 | DM-DA | 3.35 | 0.0026 |
| DA-VS | 5.32 | 6.43e-7 | SC-VS | 5.58 | 3.44e-7 |
| DA-SM | 5.38 | 6.43e-7 | SC-SM | 5.94 | 3.44e-7 |
| DA-DA | 6.93 | 7.65e-10 | SC-DA | 5.43 | 6.02e-7 |
| VA-VS | 3.33 | 0.0026 | SC-LM | 3.25 | 0.0031 |
| VA-VA | 2.86 | 0.0099 | SC-DM | 2.88 | 0.0099 |
| LM-SM | 5.33 | 6.43e-7 | SC-SC | 4.60 | 1.82e-5 |

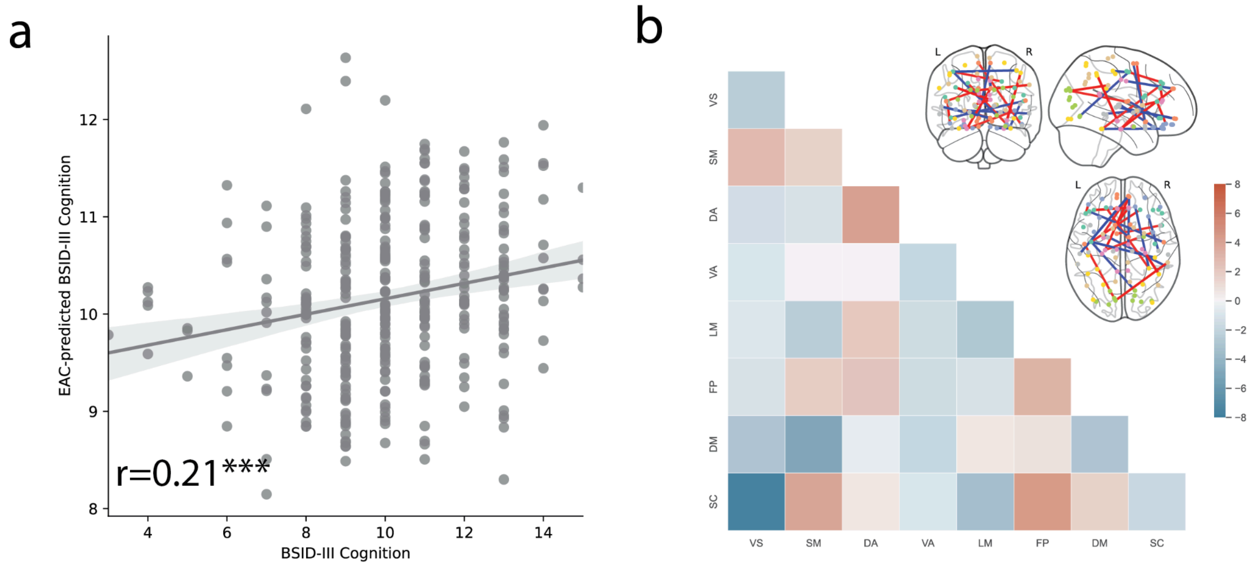


**Fig. S2. Prediction of cognition with edge controllability (a) Predicted BSID-III cognitive scores** were significantly correlated with the observed BSID-III language scores (r=0.21, p=8.28e-5)**. (b) Edge average controllability of the COG network** predicts the cognitive outcomes at 18 months of term infants, with each element showing the number of edges between each pair of canonical networks in the cognition predictive network**.**

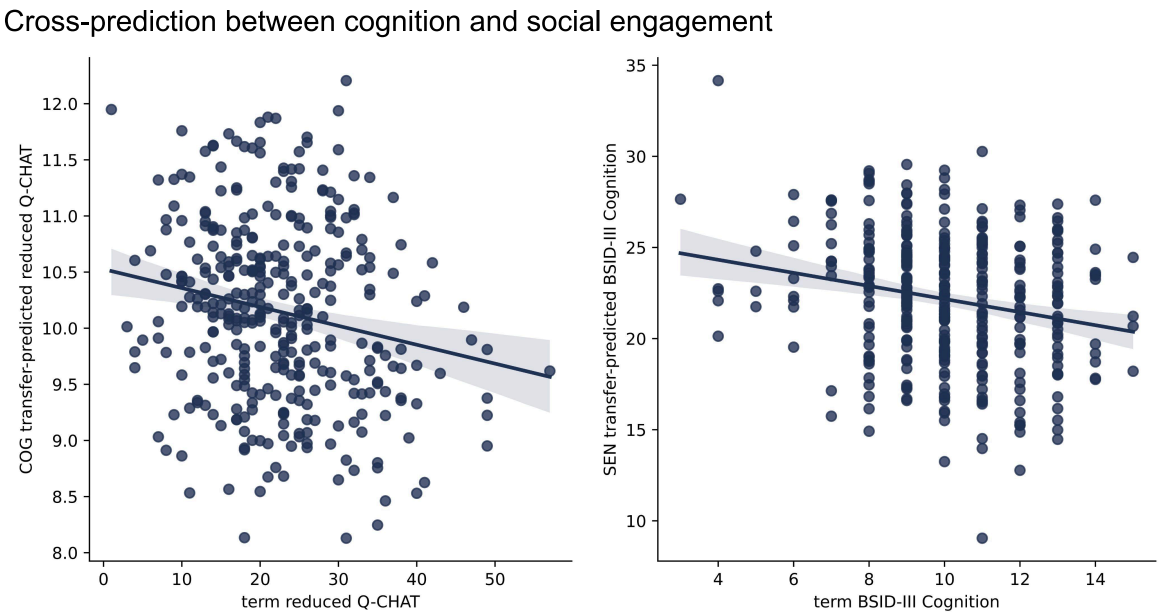


**Fig. S3. Cross-prediction between cognition and social engagement. (a)** The pre-trained SEN model predicts the cognitive outcome (r=-0.21,p=4.87e-5). **(b)** The pre-trained cognitive model predicts the social engagement outcomes (r=-0.19, p=2.96e-4).

Table S2 Quantitative Checklist of Autism in Toddlers [29]

| |
|---|
| 1. Does your child look at you when you call his/her name? |
| 2. How easy is it for you to get eye contact with your child? |
| 3. When your child is playing alone, does s/he line objects up? |
| 4. Can other people easily understand your child's speech? |
| 5. Does your child point to indicate that s/he wants something (e.g. a toy that is out of reach)? |
| 6. Does your child point to share interest with you (e.g. pointing at an interesting sight)? |
| 7. How long can your child's interest be maintained by a spinning object (e.g. washing machine, electric fan, toy car wheels)? |
| 8. How many words can your child say? |
| 9. Does your child pretend (e.g. care for dolls, talk on a toy phone)? |
| 10. Does your child follow where you're looking? |
| 11. How often does your child sniff or lick unusual objects? |
| 12. Does your child place your hand on an object when s/he wants you to use it (e.g. on a door handle when s/he wants you to open the door, on a toy when s/he wants you to activate it)? |
| 13. Does your child walk on tiptoe? |
| 14. How easy is it for your child to adapt when his/her routine changes or when things are out of their usual place? |
| 15. If you or someone else in the family is visibly upset, does your child show signs of wanting to comfort them (e.g. stroking their hair, hugging them)? |
| 16. Does your child do the same thing over and over again (e.g. running the tap, turning the light switch on and off, opening and closing doors)? |
| 17. Would you describe your child's first words as: |
| 18. Does your child echo things s/he hears (e.g. things that you say, lines from songs or movies, sounds)? |
| 19. Does your child use simple gestures (e.g. wave goodbye)? |
| 20. Does your child make unusual finger movements near his/her eyes? |
| 21. Does your child spontaneously look at your face to check your reaction when faced with something unfamiliar? |
| 22. How long can your child's interest be maintained by just one or two objects? |

| |
|---|
| 23. Does your child twiddle objects repetitively (e.g. pieces of string)? |
| 24. Does your child seem oversensitive to noise? |
| 25. Does your child stare at nothing with no apparent purpose? |
